# Ocean acidification increases susceptibility to sub-zero air temperatures in ecosystem engineers (*Mytilus* sp.): a limit to poleward range shifts

**DOI:** 10.1101/2022.06.30.498370

**Authors:** Jakob Thyrring, Colin D. MacLeod, Katie E. Marshall, Jessica Kennedy, Réjean Tremblay, Christopher D. G. Harley

## Abstract

Ongoing climate change has caused rapidly increasing temperatures, and an unprecedented decline in seawater pH, known as ocean acidification. Increasing temperatures are redistributing species towards higher and cooler latitudes which are most affected by ocean acidification. Whilst the persistence of intertidal species in cold environments is related to their capacity to resist sub-zero air temperatures, studies have never considered the interacting impacts of ocean acidification and freeze stress on species survival and distribution. A full-factorial experiment was used to study whether ocean acidification increases mortality in *Mytilus* spp. following sub-zero air temperature exposure. We examined physiological processes behind variation in freeze tolerance using ^1^H NMR metabolomics, analyses of fatty acids, and amino acid composition. We show that low pH conditions (pH = 7.5) significantly decrease freeze tolerance in both intertidal and subtidal populations of *Mytilus* spp. Under current day pH conditions (pH = 7.9), intertidal *M. trossulus* were more freeze tolerant than subtidal *M. trossulus* and *M. galloprovincialis*. Opposite, under low pH conditions, subtidal *M. trossulus* was more freeze tolerant than the other groups. We observed a marked shift from negative to positive metabolite-metabolite correlations across species under low pH conditions, but there was no evidence that the concentration of individual metabolites or amino acids affected freeze tolerance. Finally, pH-induced changes in the composition of cell membrane phospholipid fatty acids had no effect on survival. These results suggest that ocean acidification can offset the poleward expanding facilitated by warming, and that reduced freeze tolerance could result in a niche squeeze if temperatures become lethal at the equatorward edge.

## 1. Introduction

The rapid rise in atmospheric CO_2_ concentration since the industrial revolution has increased global air and water temperatures and caused ocean pH to decline (a process termed ocean acidification [OA]) at rates unprecedented in the geological history (Hönisch et al., 2012). These environmental changes are causing species range shifts and cascading ecological effects across the globe, resulting in regime shifts and altered food web structures (Kortsch et al., 2012; Wernberg et al., 2016). For example, the fish assemblage around the Svalbard archipelago, located in the Arctic Ocean (78°N), is borealizing as Arctic species have retracted northwards to cooler areas while boreal species have become dominant (Fossheim et al., 2015). Co-occurring OA is, furthermore, predicted to have severe consequences for marine organisms and communities, and a large body of research has shown a wide range of negative effects. Decreased pH weakens shell production (MacLeod and Poulin, 2015) and increases dissolution in calcifying organisms, which are therefore generally more vulnerable to OA compared to other organisms (Kroeker et al., 2010). Ocean acidification has also been found to increase heart rates in some invertebrate species (Lim and Harley, 2018) and alter benthic community structure (Brown et al., 2018). Elevated temperatures and OA have furthermore been observed to interact in various ways, causing heterogenic physiological responses across species, depending on taxon and life-stage (Harvey et al., 2013). Indeed, these two stressors may disproportionally alter species interactions and biodiversity in marine ecosystems (Franzova et al., 2019; Nagelkerken and Munday, 2016).

While the vast majority of OA and climate change research has focused on lower latitude systems, studies have rarely considered the impacts on species at their poleward edge. The poleward edge of subtidal ectotherms is determined by low water temperatures (Sunday et al., 2012), however the distribution of intertidal species is also controlled by their capacity to tolerate sub-zero air temperatures during emersion (Kennedy et al., 2020; Reid and Harley, 2021; Thyrring et al., 2019). On rocky shores, canopy-forming macroalgae shelter the understory communities from extreme sub-zero air temperatures (Sejr et al., 2021), and where cold enough, an ice foot forms on the rocky surface, creating a warmer protective microhabitat increasing survivorship of intertidal organisms residing below (Scrosati and Eckersley, 2007; Thyrring et al., 2017a). However, as temperatures increase at the northern range edge where ice forms, organisms face sub-zero air temperatures when emerged at low tides as the ice foot melts, offsetting the otherwise facilitative effect of ocean warming on range expansions.

To survive sub-zero air temperature exposure, ectothermic animals depend on various freeze tolerance mechanisms (Storey and Storey, 1996; Toxopeus and Sinclair, 2018), yet despite the principle importance of freeze tolerance in sessile intertidal species, the underlying physiological processes remain poorly understood (Kennedy et al., 2020). Suggested mechanisms behind freeze mortality are excessive osmotic stress and structural damage to cell membranes, as ice forms in the extracellular water, dehydrating the cell and destabilizing the membrane (Meryman, 1971; Storey and Storey, 1988). To avoid cell dehydration, some species accumulate cryoprotectants, such as metabolites (low molecular weight cryoprotectants), to protect against intracellular osmotic stress as water is lost to the extracellular space. In intertidal bivalves, metabolites and anaerobic byproducts such as trimethylamine n-oxide (TMAO), betaine, taurine and strombine, likely to act as cryoprotectants, increasing freeze tolerance (Kennedy et al., 2020; Loomis et al., 1988). While under-explored, it also appears that many intertidal species may have an array of ice binding proteins that help manage ice growth and propagation (Box et al., 2022).

Freeze tolerance in some ectotherms is also associated with the composition of the cell membrane phospholipid fatty acids, which are sensitive to temperature variation (Hazel, 1995). Functional membranes must exist in a fluid liquid-crystalline phase maintained by the composition of the phospholipids. Low temperatures decrease membrane fluidity, and the membrane becomes partly dysfunctional, losing selective properties and leaking cell contents (Hazel 1995). Ectotherms can counteract this effect by desaturating the membrane (increasing the proportion of unsaturated phospholipids) and adjusting cholesterol levels. This mechanism, termed homeoviscous adaptation, has been shown in a wide range of marine and terrestrial animals (Storey and Storey, 1988), and intertidal bivalves can remodel phospholipids in response to temperature changes (Pernet et al., 2007; Thyrring et al., 2017c; Williams and Somero, 1996). Despite this progress on the mechanisms of cold and freeze tolerance in intertidal species, it is completely unknown whether OA interacts with these. High latitude cold water is able to absorb significantly more CO_2_ than lower latitude warmer water, and therefore seawater pH is decreasing most rapidly at these latitudes (Fassbender et al., 2017). Ocean acidification decreases pH in osmoconformers with a low capacity to regulate internal pH levels (e.g., bivalves), which could lead to disruption of cellular processes, and shifts in osmotic balance (Wittmann and Pörtner, 2013; Zhao et al., 2020). Thus, OA may decrease freeze tolerance and increase animal vulnerability to sub-zero air temperature exposure, yet the interaction between OA and freeze tolerance interactions remains to be explored.

Bivalves of the genus *Mytilus* are distributed in intertidal habitats in both the Northern and Southern Hemisphere (Hilbish et al., 2000; Mathiesen et al., 2017). *Mytilus* sp. are commercially, and ecologically important ecosystem engineers creating habitats for a diverse associated fauna and are widely used as model organisms for studying impacts of various stressors (Barrett et al., 2022; Telesca et al., 2019; Thyrring et al., 2015). *Mytilus* sp. can survive tissue freezing, and are expanding at higher latitudes in response to global warming (Thyrring et al., 2017a), however, the performance and survival of *Mytilus* sp. at their poleward edge remain poorly understood.

The focus of this study is two *Mytilus* spp. found in British Columbia, Canada; the invasive Mediterranean mussel *M. galloprovincialis*, and the native bay mussel *M. trossulus*, allowing a comparison of responses among native and invasive species. By investigating the effects of OA on freeze tolerance in these species, we test the hypothesis that OA will generally increase mortality in intertidal species living near their poleward range edge due to an increased susceptibility to sub-zero air temperatures during emersion. Mussels from both the intertidal and subtidal realm were investigated to detect whether previous exposure to air has any effects on freeze tolerance. Specifically, we predict that (1) intertidal animals are more freeze tolerant than subtidal conspecifics, (2) native *M. trossulus* is more freeze tolerant than *M. galloprovincialis*, and (3) OA will increase freeze mortality in both species. Finally, we hypothesize that mechanistic processes including (I) a destabilized cell membrane caused by variation in the unsaturation state of membrane phospholipids, (II) and variation in the composition and concentration of selected molecular cryoprotectants will explain variation in freeze tolerance.

## 2. Materials and Methods

### 2.1 Animal collection and holding conditions

Three categories of *Mytilus* mussels were collected on 8–10 December 2019 in the strait of Georgia, British Columbia, Canada; 1) subtidal *M. galloprovincialis* obtained from an aquaculture farm at Saltspring Island, 2) subtidal *M. trossulus* collected from floating docks at the Jericho Royal Vancouver Yacht Club in the Burrard Inlet, and 3) intertidal *M. trossulus* collected at low tide from Tower Beach in the Burrard Inlet (Collection permit number XMCFR 7 2019; Fisheries and Oceans Canada). Intertidal *M. galloprovincialis* was not considered as no intertidal populations are established in region. All mussels were kept for a 72-hour adjustment period in aerated aquaria of similar environmental conditions as the collection site measured on the days of collection (7°C, pH = 7.9, and salinity 20.5). No mussels died during the adjustment period.

Prior to sub-zero air temperature exposure, mussels were maintained in low (7.50 pH) or control (7.90 pH) conditions for 10 days using three incubators (Panasonic MIR 154, Panasonic, Japan) for low pH conditions and three for control conditions (no mussels died during the 10 days). Each incubator was set to 7°C and contained three 5L glass aquarium that held three envelopes of 0.5cm gauge, rigid plastic mesh (25×24cm) that separated the three categories of mussel but allowed easy flow through of seawater (salinity 20-21). Each envelope was marked with a different colored zip tie to indicate which category of mussel it contained.

### 2.2 Seawater manipulation

The two selected pH conditions covered a realistic range of pH values currently observed or predicted for the Southern Strait of Georgia; a control (pH = 7.9) and a low pH treatment (pH = 7.5) (Ianson et al., 2016). The low pH treatment was established by using two Smart-Trak® mass flow controllers (Sierra Instruments, Inc., CA, USA) to mix 100% CO_2_ (PraxAir Canada Inc., CB, Canada) and CO_2_-free air, which was then bubbled into three acidified seawater aquaria to achieve target values of 7.50 pH. CO_2_-free air was generated by using a small compressor to pump ambient air through a 500 mL Nalgene canister that contained Soda Lime (Ormond Veterinary Supply Ltd., ON, Canada). A flow rate of 3.3 cm^3^/s of 100% CO_2_ gas and 4.11 L/min of CO_2_-free air was used to reach the target pH. Our system also removed moisture from ambient air to protect the mass flow controllers from water damage. This was achieved by running the ambient-air gas lines through a small refrigerator to reduce air temperature and cause water to precipitate into a water trap, and by installing a second 500 mL Nalgene containing desiccant (WA Hammond Drierite, OH, USA) in series with the soda lime container. Control conditions (7.90 pH) were maintained by mixing ambient air and CO_2_-free air. The use of CO_2_-free air was necessary as ambient air was artificially high in CO_2_ due to poor ventilation in the lab. As with the low pH treatment, ambient air was pumped through a Soda Lime-filled 500 mL Nalgene canister using a small compressor before being connected to a three-way splitter and bubbled into seawater aquaria, while ambient air was bubbled into the control aquaria using a second set of tubes connected to low power aquarium air pumps (Fusion 700 Air Pump). Air flow from the small aquarium pumps was fine-tuned by placing an adjustable clamp on the flexible tubing which connected pump and air stone to increase or decrease the flow of CO_2_ enriched ambient air to achieve 7.90 pH. Carbon dioxide in ambient “lab” air was monitored constantly using a Qubit S151 CO_2_ gas analyzer (Qubit Systems, ON, Canada), which showed that CO_2_ fluctuated during the day between 400 ppm and 600 ppm CO_2_ reaching the maximum during the day while people were working in the lab. Consequently, the input of ambient air into control tanks was monitored and adjusted daily (mainly during the day) to maintain target pH values. Prior to adjusting seawater pH, we mixed filtered seawater (provided by the Vancouver Aquarium and transported by the City of Vancouver) with de-chlorinated distilled fresh water to create a 20-21 ppt solution which was the salinity recorded at the collection sites. Mussels were able to feed on phytoplankton naturally occurring in the water, and we replaced 50% of the seawater from each tank daily to prevent the buildup of faeces and maintain uniform seawater chemistry parameters.

### 2.3 Carbonate chemistry

Seawater pH was measured daily in all aquaria using a hand-held pH meter (Table 1 - Oakton pH 450 (±0.01 pH), Oakton Instruments, IL, USA) calibrated with two saltwater buffers, as described in (MacLeod et al., 2015) to provide pH measurements on the Total Hydrogen Ion Scale (pH_T_). To further characterize the seawater carbonate chemistry, seawater samples (300 mL) were collected from one randomly selected aquarium in each incubator at the start and end of the experiment. These samples were fixed with a saturated solution of mercuric chloride (RICCA Chemical Company, TX, USA) and analyzed using the “burke-o-lator” at Hakai Institute (Quadra Island, BC, Canada) (for details of this system see (Evans et al., 2019)). This analysis generated values for DIC and pCO_2_ which were then used in combination with temperature and salinity data to calculate all relevant carbonate parameters (Supplementary Table S1) using the MATLAB version of CO_2_SYS (van Heuven et al., 2011).

**Table 1:**
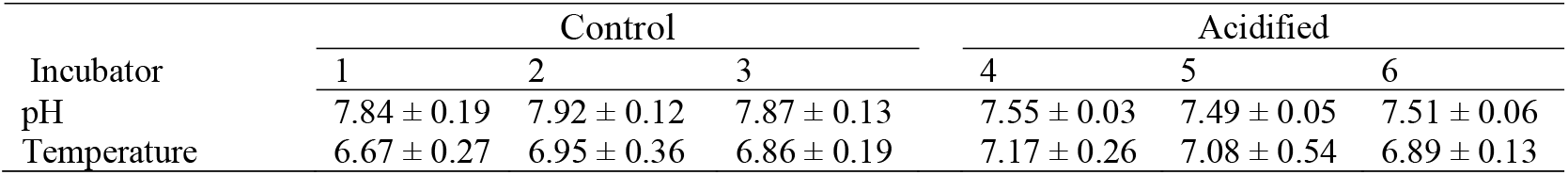
Mean (± standard deviation) pHT and temperature measured directly in aquaria in each incubator.

### 2.4 Sub-zero temperature exposure

After 10 days, mussels were exposed to seven sub-zero air temperatures (−5, −6, −7, −8, −9, −12, −15 °C) for two hours by placing animals in individual plastic tubes inserted in wells drilled into a precooled aluminum block cooled by refrigerated circulation baths (Thermo Fisher Scientific Inc., MA, USA). Fifteen mussels (mean shell length 37.69 mm ± 3.14 s.d.) from each mussel category (subtidal *M. galloprovincialis* and *M. trossulus*, and intertidal *M. trossulus*) and pH condition (pH =7.9 and pH = 7.5) were used at every temperature for a total of 720 mussels. Individual body temperatures were recorded at 0.5 s intervals using Type-T thermocouples (Omega, QC, Canada) placed next to the shell inside the plastic tube and connected to TC-08 thermocouples interfaces (Pico Technology, United Kingdom) that interfaced to a computer running PICOLOG software (Picotech, United Kingdom), which continuously monitored body temperatures. Continuously body temperature monitoring allowed us to determine any exothermic release of heat owing to ice formation. The lowest temperature prior to this event is termed the supercooling point (SCP), and the SCP indicates that internal ice formation occurred. After two hours of sub-zero air exposure, all mussels were transferred back to their respective pre-freezing pH condition aquaria for recovery where they were monitored daily for five days to record mortality. Mortality was checked daily with mussels considered dead if they did not close their shells when touched. Dead mussels were immediately removed from the aquaria, and had their shell length measured to nearest mm.

### 2.5 Amino acid analysis

Total amino acid analysis was performed on gill tissue from mussels collected after 10 days of pH exposure (mean dry weight = 15.46 mg ±0.61 s.d., *n* = 5) at the Proteomics, Analytics, Robotics & Chemical Biology Centre (SPARC - https://lab.research.sickkids.ca/sparc-molecular-analysis/services/amino-acid-analysis/), Hospital for Sick Children, Toronto, ON, Canada, using the Water Pico-Tag System (Water Corporation, WA, USA). The final concentration of each amino acid was calculated in μg·mg^−1^ and then expressed as relative concentration (% of total amino acids). It should be noted that this amino acid analysis did not allow discrimination between Asn/Asp and Gln/Glu.

### 2.6 Fatty acid analysis

Fatty acid (FA) analyses were performed on gills (collected after 10 days of pH exposure), a tissue exposed directly to temperature changes (mean wet weight = 0.38g ±0.1 s.d., *n* = 5). Total lipids were extracted by grinding in a dichloromethane: methanol (2:1, v/v) solution following a slightly modified Folch procedure (Parrish, 1999). Lipid extracts were separated into neutral and polar fractions by column chromatography on silica gel micro-columns (30 × 5 mm i.d., packed with Kieselgel 60, 70-230 mesh; Merck, Germany) using chloroform: methanol (98:2, v/v) to elute neutral lipids, followed by methanol to elute polar lipids (Marty et al., 1992). Fatty acid profiles were determined on fatty acid methyl esters (FAMEs) using sulphuric acid:methanol (2:98, v/v) and toluene. FAMEs of neutral and polar fractions were concentrated in hexane, and the neutral fraction was purified on an activated silica gel with 1 ml of hexane:ethyl acetate (1:1 v/v) to eliminate free sterols. FAMEs were analyzed in the full scan mode (ionic range: 60-650 m/z) on a Polaris Q ion trap coupled multi-channel gas chromatograph ‘Trace GC ultra ’(Thermo Scientific, USA) equipped with an auto sampler model Triplus, a PTV injector and a mass detector model ITQ900 (Thermo Scientific, USA). The separation was performed with an Omegawax 250 (Supelco) capillary column with high-purity helium as a carrier gas. Data were treated using Xcalibur v.2.1 software (Thermo Scientific, USA). Methyl nondecanoate (19:0) was used as an internal standard. FAMEs were identified and quantified using known standards (Supelco 37 Component FAME Mix and menhaden oil; Supleco) and were further confirmed by mass spectrometry (Xcalibur v.2.1 software).

### 2.7 ^1^H NMR analysis

One-dimensional, 600 MHz proton nuclear magnetic resonance spectroscopy (^1^H NMR) was used to measure the metabolite profiles of gill tissue (collected after 10 days of pH exposure). ^1^H NMR is ideal for measuring low molecular weight, polar metabolites such as osmolytes and anaerobic byproducts. Sample preparation was based on (Cappello et al., 2013). A 100 mg sample of gill tissue was excised (*n* = 5), dried with a Kimwipe to remove excess water and frozen at −80°C. Frozen tissue was homogenized in 400 μl cold methanol and 85 μl cold water-xylitol solution (5 mM xylitol as an internal control) using a bead homogenizer (Bullet Blender 50 Gold Model: BBX24, Next Advance) with approximately 200 μl of 3.2 mm round stainless steel beads, for 10 min at setting 8 in 1.5 ml microcentrifuge vials. After adding 400 μl chloroform and 200 μl water to the samples, they were vortexed for 60 s, left on ice for 10 min for phase separation, and centrifuged for 5 min at 2000 rpm. The upper methanol layer (600 μl) containing the polar metabolites was transferred into new vials, dried in a centrifugal vacuum concentrator (Eppendorf 5301), and then stored at −80°C. Immediately prior to ^1^H NMR analysis, the dried polar extracts were resuspended in 600 μl of 0.1 mol/l sodium phosphate buffer (pH 7.0, 50% deuterium oxide, Sigma-Aldrich) containing 1 mmol/l 2,2-dimethyl-2-sila-pentane-5-sulfonate (DSS; Sigma-Aldrich) as internal reference. The mixture was vortexed for 60 s and transferred to a 5 mm NMR tube.

^1^H NMR spectra were acquired using Bruker Avance 600 with cryoprobe and Bruker Avance III 600 spectrometers. TopSpin software version 2.1 (Bruker) was used to process spectra collected with the Bruker Avance 600 spectrometer with cryoprobe, and TopSpin version 3.5 (Bruker) was used with the Bruker Avance III 600 spectrometer. Experiments required 15 minutes of acquisition time and were performed at room temperature.

Peak identification of the NMR spectra was performed with Chenomx NMR Suite 9.0 (Chenomx, AB, Canada) that uses the Human Metabolome Database compound spectral reference library (Wishart et al., 2018). First, line broadening of 2.5 Hz, automatic phase correction, and manual baseline correction were performed with Chenomx Processor (within the Chenomx NMR Suite software). Then, determination of metabolite concentrations was performed using Chenomx Profiler, which determines the concentrations of individual metabolites using the concentration of a known DSS signal. Metabolite concentrations are reported as mmol/100 mg gill wet mass.

### 2.8 Statistical analysis

#### 2.8.1 Survival

Statistical analyses were performed using the R software (R version 3.5.2). A logistic regression model was used to calculate LLT_50_ values (the lower lethal temperature where 50% of the population survived). A binomial generalized linear model (GLM) with a logit link function, was used to determine the effects of air temperature and pH treatment on survival within each mussel category, and the difference in LLT_50_ were estimated using 95% confidence intervals (CI) with non-overlapping CI indicating a significant difference (Deere et al., 2006). Differences in the supercooling point (SCP) among mussel categories and pH treatment was analyzed using a two-way ANOVA. Final models were validated by plotting residuals versus fitted values, versus each covariate in the model (Zuur and Ieno, 2016).

#### 2.8.2 Metabolomics and fatty acids

Generalized linear models and ANOVAs were used to determine which metabolites and fatty acids differed significantly after low pH exposure. Pearson’s correlations were furthermore used to detect alternations in metabolite-metabolite associations following low pH conditions (Jahagirdar and Saccenti, 2020), and relationships were visualized using heat maps. The fatty acids explaining most of the dissimilarity between mussel categories and pH treatments were identified using a SIMPER analysis (See the full list of FA founds in Supplementary Table S2). This analysis revealed that 13 fatty acids explained ~90% of the Bray–Curtis dissimilarity amongst fatty acid profiles between the control and low pH environment (Table 2). We therefore focused all subsequent fatty acid analyses on these 13 fatty acids. Principle component analysis (PCA) was used to the interpretation of differences in the metabolomic composition among mussel categories. ANOVAs were used to evaluate differences in the concentrations of amino acids, and GLMs to evaluate the distribution of saturated fatty acids (SFA), monounsaturated fatty acids (MUFAs), polyunsaturated fatty acids (PUFAs), and the unsaturation index (UI), which is an index for the number of double bonds per 100 molecules of fatty acids, among the three mussel categories and pH Treatment. Post-hoc pair-wise tests were used to compare significant treatment effects (P <0.05). Detailed data exploration was carried out prior to any analysis (Zuur et al., 2010). Once valid models were identified, we re-examined the residuals to ensure all model assumptions were acceptable.

**Table 2:** Mean percentage (± standard deviation) of 13 fatty acids contributing ~90% of the differences in fatty acid composition after subjection to control (pH = 7.9) and acidified (pH = 7.5) conditions (n = 5). UI = unsaturation index. Bold numbers indicate significant differences among the control and low pH treatment within a mussel category (p < 0.05).

| Fatty acid | Intertidal - *M. trossulus* |  | Subtidal - *M. trossulus* |  | Subtidal - *M. galloprovincialis* |  |
| --- | --- | --- | --- | --- | --- | --- |
|  | Acidified | Control | Acidified | Control | Acidified | Control |
| 16:0 | 9.53±0.89 | 8.72±1.21 | 16.56±6.62 | 10.21±1.21 | 12.95±5.17 | 8.83 ±1.44 |
| 18:0 | 2.57±0.34 | 2.53±0.17 | 4.30±1.98 | 2.28±0.38 | 3.38±1.26 | 2.53±0.67 |
| **16:1ω7** | 1.01±0.1 | 1.2±0.22 | **1.58±0.33** | **0.97±0.19** | **1.71±0.33** | **1.07±0.27** |
| 17:1ω7 | 1.71±0.63 | 4.07±4.16 | 1.94±1.14 | 1.43±1.1 | 2.26±1.63 | 2.04±2.15 |
| **18:1ω7** | 1.48±0.12 | 1.37±0.37 | **1.91±0.52** | **1.21±0.2** | 2.08±0.58 | 1.5±0.34 |
| **20:1ω9** | 4.31±0.3 | 4.12±0.46 | **6.21±1.13** | **4.86±0.61** | 5.43±1.32 | 4.37±0.48 |
| 18:2ω6 | 0.91±0.1 | 0.94±0.35 | 1.46±0.67 | 1.31±0.25 | 0.79±0.36 | 0.64±0.14 |
| 18:3ω3 | 1.12±0.07 | 1.21±0.5 | 1.04±0.57 | 1.52±0.19 | 0.31±0.1 | 0.36±0.09 |
| **20:2** | 9.41±0.76 | 8.82±1.05 | **11.96±1.21** | **9.87±0.58** | 9.78±1.79 | 8.67±1.41 |
| 20:4ω6 | 9.70±1.1 | 9.81±0.9 | 3.76±2.61 | 6.07±0.76 | 6.95±2.43 | 9.7±1.37 |
| **20:5ω3** | 9.78±1.15 | 10.74±1.97 | **4.87±3.59** | **12.19±1.05** | 7.0±2.98 | 10.21±1.68 |
| **22:2-NMI** | 11.54±0.84 | 10.86±1.5 | **12.72±0.95** | **9.55±0.66** | 13.53±1.5 | 12.19±1.14 |
| **22:6ω3** | 14.13±0.99 | 13.73±0.69 | **6.36±4.86** | **16.22±0.48** | **9.7±4.61** | **15.88±2.06** |
| **UI** | 234.73±6.14 | 238.84±8.75 | **150.94±55.19** | **244.97±4.59** | **187.59±46.35** | **242.92±8.78** |

## 3. Results

### 3.1 Survival

Validation of ANOVAs and GLM models indicated no violation of model assumptions. There were no significant effects of pH conditions (ANOVA; p > 0.05) or mussel categories (ANOVA; p > 0.05 on the SCP, and freezing of the tissue was observed in all mussels exposed to temperatures below −7°C.

We investigated the effect of pH and sub-zero air temperature on survival using GLMs. There were no significant interactions between the effect of pH and air temperature on any mussel category, and the interaction term was excluded in the final GLM models. Lower sub-zero air temperature significantly decreased the survival of mussel in all three categories exposed to both control and low pH conditions (Fig. 1; Supplementary Table S3). Under control conditions (pH = 7.9), the lower lethal temperature at which 50% of the population perish (LLT_50_) was significantly lower in intertidal *M. trossulus* (−10.56 °C ± 0.80 CI) compared to subtidal *M. trossulus* (−9.12°C ± 0.48 CI) and subtidal *M. galloprovincialis* (−7.62 °C ± 0.49 CI), which was the least freeze tolerant species (Fig. 1). Following exposure to low pH (pH = 7.5), survival significantly decreased after sub-zero air exposure in all three mussel categories (Fig. 1; Supplementary Table S3). Accordingly, the LLT_50_ of intertidal *M. trossulus* was −7.53°C ± 0.26 CI, while the LLT_50_ was −8.04°C ± 0.32 CI and −6.69°C ± 0.17 CI for subtidal *M. trossulus* and subtidal *M. galloprovincialis*, respectively. Thus, subtidal *M. trossulus* was the most freeze tolerant category under low pH conditions. It should be noted that only one *M. galloprovincialis* (6.66%) survived exposure to −8°C.

**Figure 1:**
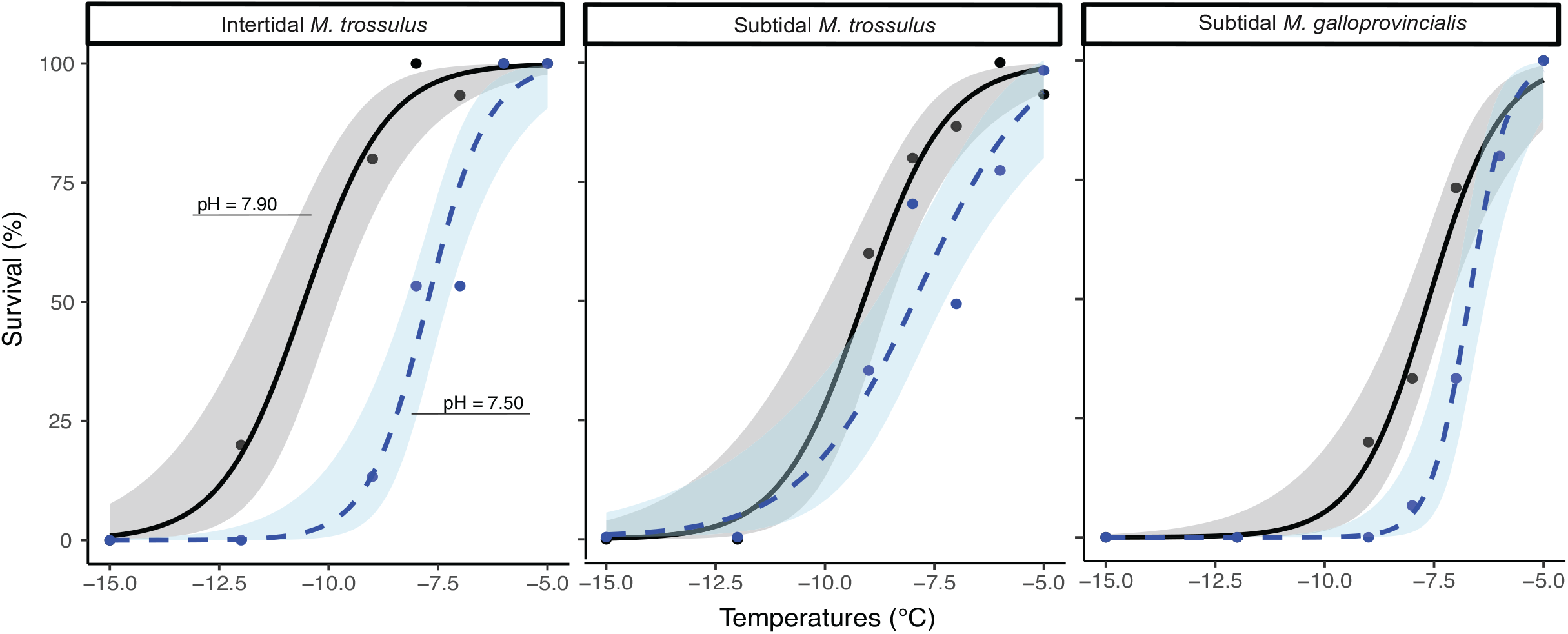
Proportion survival in intertidal *Mytilus trossulus*, subtidal *M. trossulus* and subtidal *M. galloprovincialis* after subjection to two pH treatments for 10 days and seven sub-zero air temperatures. Lines indicate fitted logistic regression models; Solid black line represent control conditions (pH =7.9) and dashed blue line represent acidified conditions (pH =7.5). Dots represent actual survival and shades areas indicate 95% confidence intervals of the fitted model.

### 3.2 Metabolomics and fatty acids

We compared the composition of metabolites using PCA plots, which showed that the three mussels categories clustered together, suggesting no differentiation in their metabolic profiles (Supplementary Fig. S1). The predominate osmolytes were alanine, aspartate, betaine, glycine, malonate, taurine, trimethylamine and trimethylamine n-oxide (TMAO) (see Supplementary Table S4 for full list of all metabolites obtained from the ^1^H NMR analysis), but we detected no significant changes in their concentration among the two pH treatments or mussel categories (Fig. 2). Likewise, were there no significant differences in the concentration of any amino acids among mussel categories or pH treatment (Table 3). A metabolite-metabolite Pearson correlation analysis of the predominate osmolytes revealed that each mussel category had a unique metabolite-metabolite relationship under control pH conditions (Fig. 3 Upper panel). Low pH conditions changed the metabolite-metabolite relationship in all three mussel categories (Fig. 3 Lower panel); A marked shift from negative to positive correlations was observed across all metabolites in *M. trossulus* where the number of positive correlations increased by circa 53% in intertidal *M. trossulus* and 109% in subtidal *M. trossulus*. In *M. galloprovincialis*, the number of positive metabolite-metabolite correlations decreased from 11 to 10 (Fig. 3).

**Figure 2:**
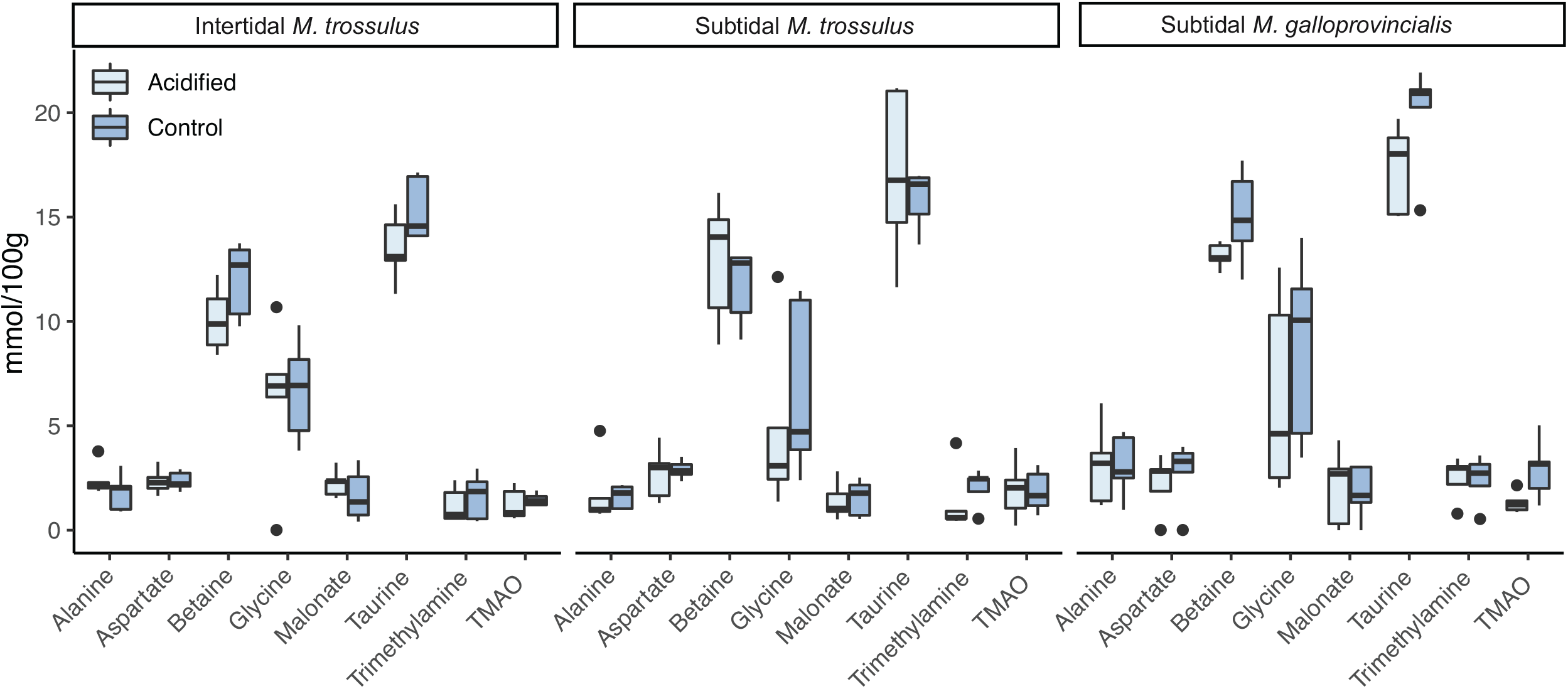
The concentration of common gill tissue osmolytes in intertidal *Mytilus trossulus*, subtidal *M. trossulus* and subtidal *M. galloprovincialis* after subjection to control (pH = 7.9) and acidified (pH = 7.5) conditions (n = 5). The horizontal line in each boxplot is the median, the boxes define the hinges (25–75% quartile) and the whisker is 1.5 times the hinges. Black dots represent data outside this range.

**Table 3:**
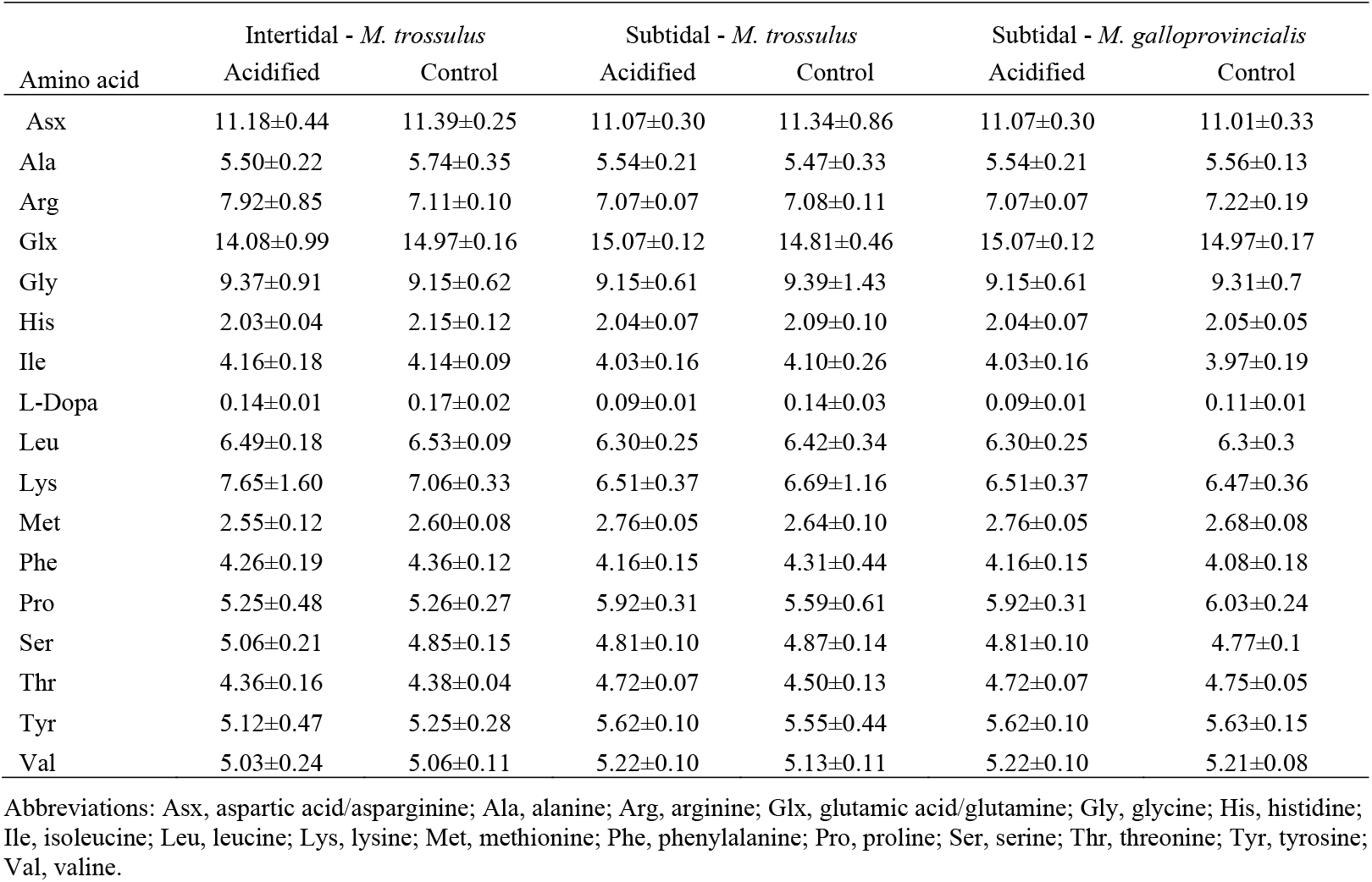
Mean (± standard deviation) amino acid composition (% total amino acid content) of gill tissue in intertidal *Mytilus trossulus*, subtidal *M. trossulus* and subtidal *M. galloprovincialis* after subjection to control (pH = 7.9) and acidified (pH = 7.5) conditions (n = 5)

**Figure 3:**
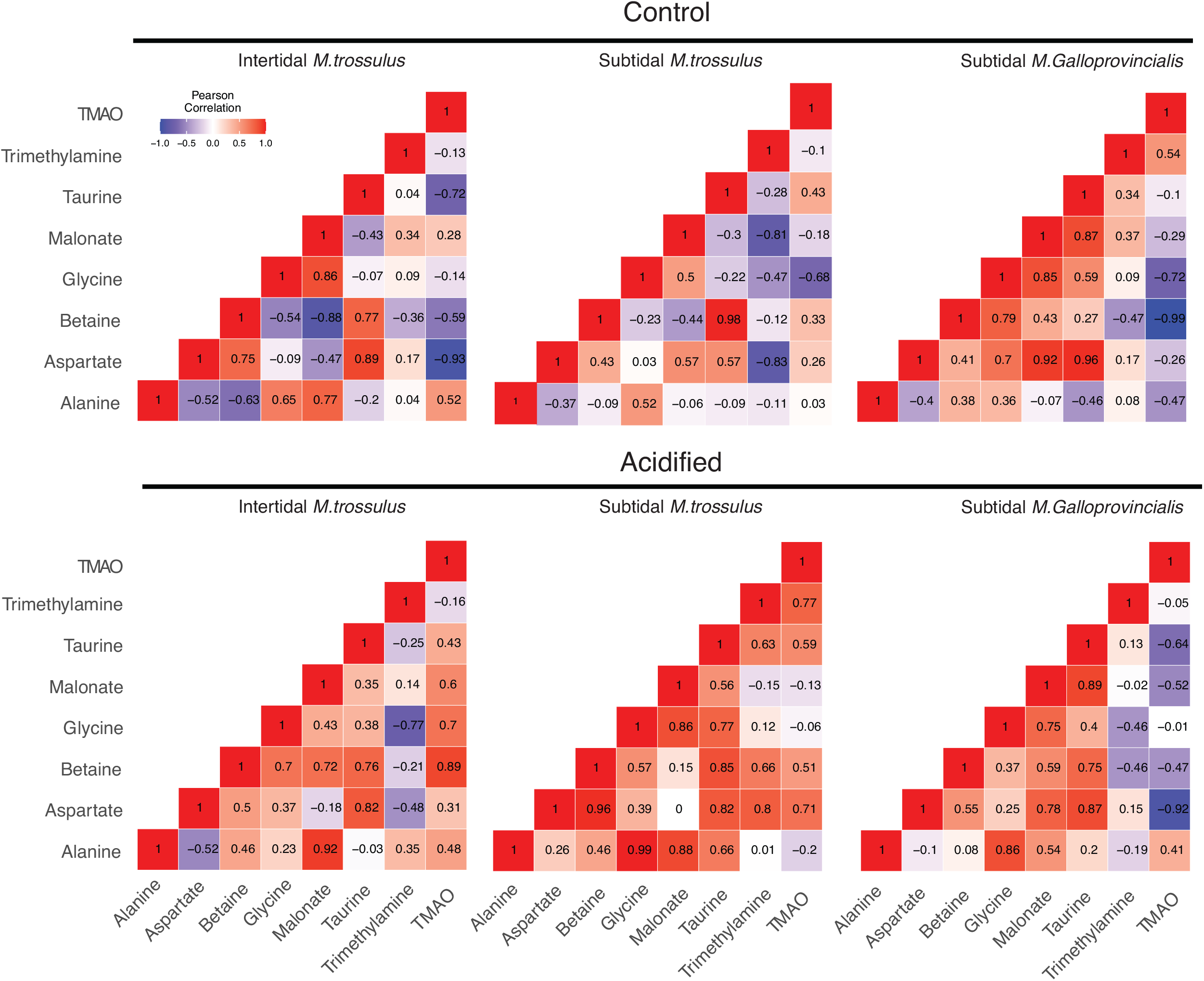
Metabolite-metabolite correlation analysis of gill tissue in intertidal *Mytilus trossulus*, subtidal *M. trossulus* and subtidal *M. galloprovincialis*. Upper panel: metabolite associations after subjection to the control (pH = 7.9) conditions. Lower panel: metabolite associations after subjection to the acidified (pH = 7.5) conditions. Positive correlations are shown in blue; negative correlations are shown in red.

Thirteen fatty acids contributed ~90% of the difference in membrane composition between the control and low pH treatment mussels (Table 2). While the fatty acid profiles in intertidal *M. trossulus* were unaffected by low pH exposure (Table 2; Fig. 4), the low pH treatment caused an increase in the amount of SFA and a decrease of PUFA in subtidal *M. galloprovincialis* and *M. trossulus* (Fig. 4), which resulted in a significant decrease in the degree of unsaturation (GLMs; p < 0.05; Table 2; Fig. 4). Accordingly, the unsaturation index (UI) was significantly higher in intertidal than subtidal *M. trossulus* after pH exposure (TukeyHSD, p = 0.0002).

**Figure 4:**
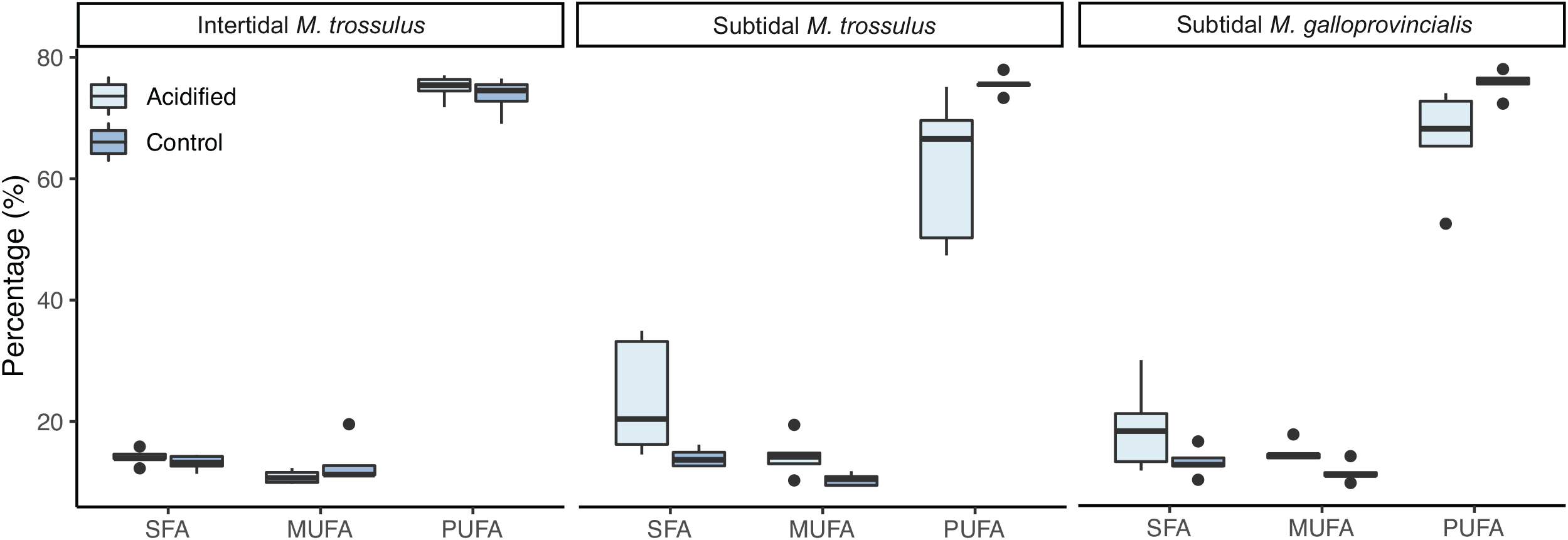
The molar percentage of saturated fatty acids (SFA), monounsaturated fatty acids (MUFA), polyunsaturated fatty acids (PUFA) of gill tissue in intertidal *Mytilus trossulus*, subtidal *M. trossulus* and subtidal *M. galloprovincialis* after subjection to control (pH = 7.9) and acidified (pH = 7.5) conditions (n = 5). The horizontal line in each boxplot is the median, the boxes define the hinges (25–75% quartile) and the whisker is 1.5 times the hinges. Black dots represent data outside this range.

### 3.3 Seawater chemistry

Mean pH measurements from the hand-held pH meter showed relatively good agreement with our target acidified treatment (7.5 pH_T_ ± 0.03-0.06 s.d.), although there was a greater degree of variability in the control treatments (7.9 pH_T_ ± 0.12-0.19 s.d.) due to fluctuating ambient pCO_2_ (Table 1; Supplementary Fig. S2). There was also some disagreement between our measured pH_T_ and pH calculated using DIC and pCO_2_ data (Supplementary Table S5). The discrepancies in our control treatments were the result of the highly variable ambient pCO_2_ and the corresponding adjustments we frequently made to our gas delivery system. On average, these fluctuations did not cause significant deviations from our target pH values, as shown by our hand-held pHT data (Table 1), but were more pronounced in the data taken from discrete, single time-point water samples. In all but one instance, the difference between directly measured and calculated pH for a single time point, i.e. the direct pHT measurement made at the time the discrete water sample was taken, was within the standard deviation of mean pH measurements taken over the two-week period. The regulation of OA simulation systems with potentiometric pH meters has been shown to be reliable (MacLeod et al., 2015), and therefore it is likely that the discrepancy between discrete and hand-held pHT data was not indicative of substantial deviations in seawater chemistry target values.

The addition of CO_2_ free air to the controls also resulted in lower-than-expected pCO_2_ values in both start and end point data from those treatments (Supplementary Table S1 and S5). We also observed some anomalous values for end point total alkalinity and DIC in our control treatment which were attributed to shell calcification and insufficient water replacement rates (Supplementary Table S1 and S5). These values were not indicative of the seawater chemistry parameters over the entire experimental period, as described above, but are included for completeness.

### 3.4 Seawater chemistry variability

In contrast to open oceans where pH is stable (Hofmann et al., 2011), daily and seasonal fluctuations can exceed 0.7 pH units in coastal ecosystems (Baumann et al., 2015; Hofmann et al., 2011; Menéndez et al., 2001; Santos et al., 2011; Semesi et al., 2009), where dense blue mussel beds have been found in areas characterized by Ω_arag_ < 0.5 (Duarte et al., 2020). Specifically, along the intertidal rocky shoreline of the Northwest Pacific, where mussels for this study was collected, pH values naturally decline below 7.6 (Ianson et al., 2016; Kroeker et al., 2016). Thus, while pH conditions in our aquariums fluctuated (control 7.84±0.19–7.92±0.12| acidified 7.49±0.05–7.55±0.03), the variability was within the range of *in situ* fluctuation rates. Our control conditions therefore represent actual *in situ* conditions, and the final average difference in pH between the control (7.88) and acidified (7.52) represented our target values (7.9 pH and 7.5 pH). The natural variation in coastal pH also challenges the common belief that Ω_arag_ should be >1 (non-corrosive conditions) to represent control conditions in OA experiments. Instead, we argue that control conditions should reflect actual pH levels on the site of collections, regardless of the Ω_arag_ level. While we acknowledge that our regulation of seawater chemistry could be improved, we believe that changes in mussel survival were caused by changes in average pH, rather than variation in pH or other parameters. Our rationale is supported by the fact that intertidal mussels exhibited the largest decrease in survival upon exposure to reduced pH plus freezing, and as they are typically exposed to much greater variability in seawater chemistry and temperature than subtidal mussel populations, it is highly unlikely that variability had the most pronounced and negative affect on this group.

## Discussion

Climate change is redistributing species towards cooler environments but understanding how different drivers interact to shape species distribution ranges is essential for predicting patterns and rates of change. At higher latitudes, expanding species face a suite of novel abiotic conditions including low temperatures, and a decreasing seawater pH (Fassbender et al., 2017). The goals of this study were to investigate the combined effect of low seawater pH and sub-zero air temperature stress on survival of two *Mytilus* spp. and compare the responses between a native and invasive congener. Intertidal individuals of the native bay mussel *M. trossulus* were significantly more freeze tolerant than subtidal *M. trossulus* individuals which were in turn more freeze tolerant than the invasive Mediterranean mussel *M. galloprovincialis*. Following exposure to acidified seawater, our data demonstrated a significant negative effect on freeze tolerance and survival across all species-habitat combinations. Interestingly, the intertidal population of *M. trossulus* was most impacted by acidification, while subtidal *M. trossulus* was the least affected, becoming most freeze tolerant. Cellular accumulation of metabolites and reconfiguration of membrane fatty acids were uncorrelated with the observed variation in survival among mussel categories under both control and acidified conditions, which could be related to short term exposure. The homeoviscous adaptation related to the inverse relationship between the unsaturation index (UI) and acclimation temperature was mainly related to 22:6ω3 and 20:5ω3 levels (Pernet et al., 2007), thus the more than two time lower content of 22:6ω3 and 20:5ω3 in subtidal *M. trossulus* in low pH condition, could suggest that long term exposure to low pH decrease the cold acclimation capacity for this group of mussels. Although, we were unable to explain the observed variation in survival, we show that exposure to acidified water changed metabolite-metabolite associations in all three mussel categories, indicating that perturbations in seawater chemistry induce molecular alterations in these species.

Under present-day conditions (our control pH treatment), both the intertidal and subtidal *M. trossulus* category of the native *M. trossulus* were more freeze tolerant than the invasive *M. galloprovincialis*. This corresponds to their geographic distribution where *M. trossulus* predominantly inhabit shorelines at higher latitude where winter sub-zero air temperatures are common, while *M. galloprovincialis* dominate on warmer low latitude shores (Hilbish et al., 2000). However, the physiological processes behind inter- and intraspecific variation in freeze tolerance remains poorly understood. In *Mytilus* spp., the accumulation of intracellular low molecular weight osmolytes increases freeze tolerance (Kennedy et al., 2020, Williams, 1970), but the accumulation of these putative cryoprotectants can only partly explain survival after sub-zero temperature exposure. For example, although individuals of *M. trossulus* living in the upper intertidal zone are more freeze tolerant than individuals from the low zone, a recent study found no differences in the concentration of metabolites among the shore levels (Kennedy et al., 2020), and no differences in the concentration of cryoprotectants were observed among our three mussel categories, despite large variation in freeze tolerance. Likewise, after ten days of exposure to acidified water that significantly reduced freeze tolerance in all three mussel categories, with the survival of the intertidal population most affected, the low pH exposure had no effect on metabolite concentration in any mussel category, offering no explanation for the observed decrease in freeze tolerance. The fact that the intertidal category was most affected by low pH support our notion that decreased survival was caused by changes in average pH, and not pH variation (see result section 3.4) because animals from more unstable environments (i.e. the intertidal) are generally more resilient to changing environmental conditions (Clark et al., 2018).

Another proposed driver of freeze tolerance is the composition of the membrane’s phospholipid where a positive relationship between survival and membrane unsaturation state (i.e., higher number of double bonds in the membrane) has been shown in some species (Bindesbøl et al., 2005; Slotsbo et al., 2016). We hypothesized that freeze tolerance in *Mytilus* mussels would also be correlated to unsaturation state, but we observed no significant differences in UI among mussel categories under control pH conditions, and our data show that PUFA and UI in subtidal *M. trossulus* and *M. galloprovincialis* decreased in response to OA, yet they were the least affected in terms of freeze tolerance. Meanwhile, intertidal *M. trossulus* had the highest unsaturation stage, but the lowest survival. The phospholipid membrane composition changes have been regarded primarily as an adaptation related to seasonal temperature variability and at our knowledge only one study showed hourly membrane lipids restructuration related to temperature for a cyprinid fish (Carey and Hazel, 1989). Our results support that phospholipid composition is of limited importance for freeze tolerance in *Mytilus* mussels, while membrane reconfiguration seems to be important for keeping membranes functional in cold water environments (Pernet et al., 2007; Thyrring et al., 2017c), thus membrane reconfiguration may be important for species to inhabit cold subtidal environments.

While we were unable to explain the variation in freeze tolerance under present-day and acidified conditions, variation in freeze tolerance among populations and congeners may be explained by high molecular weight cryoprotectants, e.g., ice binding proteins, not measured here. Indeed, the influence of antifreeze proteins on freeze tolerance in *Mytilus* ought to be explored further as their potential role seems to vary among populations (Box et al., 2022; Loomis, 1995). Furthermore, thermal tolerance variation may be explained at the gene level (Clark et al., 2021; Peck et al., 2015). A recent study highlighted that differences in the expression of heat shock genes and aquaporins plays a central role in determining freeze tolerance in northern barnacles species (Marshall et al., 2018), and heat shock proteins (HSPs) have been linked to sub-zero temperature survival in insects (Rinehart et al., 2007). Populations from variable environments (such as the intertidal zone or polar regions) are regularly exposed to unpredictable conditions, which can introduce a front-loading of stress genes that enable individuals to better cope with unfavorable conditions. For example, exposure to variable temperatures increases the overall thermal tolerance in the limpet *Lottia digitalis* (Drake et al., 2017), and studies have revealed animals from benign static environments are more vulnerable to unpredictable thermal stress (Marshall et al., 2021; Wang et al., 2020). Front-loading of genes is known from other marine species (Clark et al., 2008; Drake et al., 2017), and freeze tolerant *Mytilus* populations may also have front-loaded genes (e.g. HSPs, aquaporins) that are constantly at a higher expression level, which transfer into resilience through faster production of stress mediating proteins (Barshis et al., 2013). Thus, since the intertidal population of *M. trossulus* are used to daily air exposure, compared to subtidal *M. trossulus* and *M. galloprovincialis*, constantly increased gene expressions may provide an explanation to the difference observed in survival following sub-zero air exposure. Similarly, the subtidal *M. trossulus* mussels were collected in the Burrard Inlet, Vancouver, where the abiotic conditions are variable (Marshall et al., 2021). The exposure to these fluctuating abiotic conditions may also trigger a front-loading of relevant stress genes, and activate the cellular stress response systems (i.e. activating stress proteins such as HSPs (Kültz, 2020)), pre-increasing tolerance to low pH exposure in subtidal *M. trossulus. Mytilus galloprovincialis* is known to express the stress regulating gene HSPA12 in response to stressful conditions (You et al., 2013). HSPA12 belongs to the HSP70 family, and through a process of gene duplication, HSPA12 acquired subtly different functions, collectively working as a stress regulator (Clark et al., 2021). These stress regulators could, among other things, explain the limited effect of OA on freeze tolerance, and generally the capacity of *M. galloprovincialis* to establish in novel environments. The expression of HSPA12 as a thermal stress mediator was also recently demonstrated in *M. edulis* (Clark et al., 2021), and it is likely that *M. trossulus* also express this protective gene family. Still, intertidal *M. trossulus* may be the most affected by OA because intertidal species generally already live close to their physiological limits and have a limited capacity to adapt to new conditions. The large increase in mortality following low pH exposure may indicate that accommodating this additional environmental stressor exceeds their physiological ability to cope with external stressors. A molecular investigation could reveal the molecular processes behind variation in freeze tolerance among populations and species, and investigations into the underlying genetic mechanisms accounting for our observations would be interesting for future research.

Overall, *Mytilus* sp. are excellent at adapting to local environments (Riginos and Cunningham, 2005), making them highly stress tolerant and capable of enduring large ranges of salinities and temperatures (Barrett et al., 2022; Nielsen et al., 2021). Combined with a long pelagic larval period, they have a strong potential to invade new regions (Cárdenas et al., 2020; Thyrring et al., 2017b). This is the first study to investigate the effect of OA on freeze tolerance, an important trait necessary for a poleward shift in the intertidal zone. While intertidal *M. trossulus* populations are found as far north as northern Greenland (Mathiesen et al., 2017), the northern distribution limit of the invasive *M. galloprovincialis* in the Northwest Pacific is set around Canada. At their range edge in the waters of British Columbia, Canada, subtidal populations face intense predation by seastars, excluding mussels from the subtidal and low intertidal (Harley, 2011). The near future pH scenario tested here (pH = 7.5) revealed that acidification weakens freeze tolerance across *Mytilus* spp. (LLT_50_ ~ 7.38 – 7.53°C in intertidal *M. trossulus* and *M. galloprovincialis*), and since winter low tides predominantly occurs at nighttime in the Northwest Pacific, occasionally exposing sessile intertidal organisms to air temperatures down to −10°C (Kennedy et al., 2020), significant annual freeze mortality events could occur in both species inhabiting the intertidal if pH continues to decline. Thus, an ongoing poleward expansion in the intertidal (where predation is less intense) could be hindered, offsetting the poleward expanding facilitated by warmer waters. Consequently, if temperatures become too high for survival at a species equatorward edge, the combined effects of predation and limited freeze tolerance could result in a niche squeeze (rather than an expansion), substantially threatening persistence of this species in some regions.

## Data availability

All data necessary for reproducing this work is freely available from the Borealis dataverse Repository (https://borealisdata.ca; DOI available upon acceptance).

## Acknowledgements

We would like to acknowledge the technical assistance and expertise of researchers at the Hakai Institute, Quadra Island, British Columbia, who conducted the chemical analysis of our discrete seawater samples, and Vancouver Aquarium and by the City of Vancouver for providing water and transportation. This research has been supported by a Marie Sklodowska-Curie Individual Fellowship (IF) under contract number 797387, and by the Independent Research Fund Denmark (Danmarks Frie Forskningsfond) (DFF-International Postdoc; case no. 7027-00060B). The funding sources was not involved in designing the study design, in the collection, analysis and interpretation of data, in the writing of the report, or in the decision to submit the article for publication.

---

**Figure S1:**
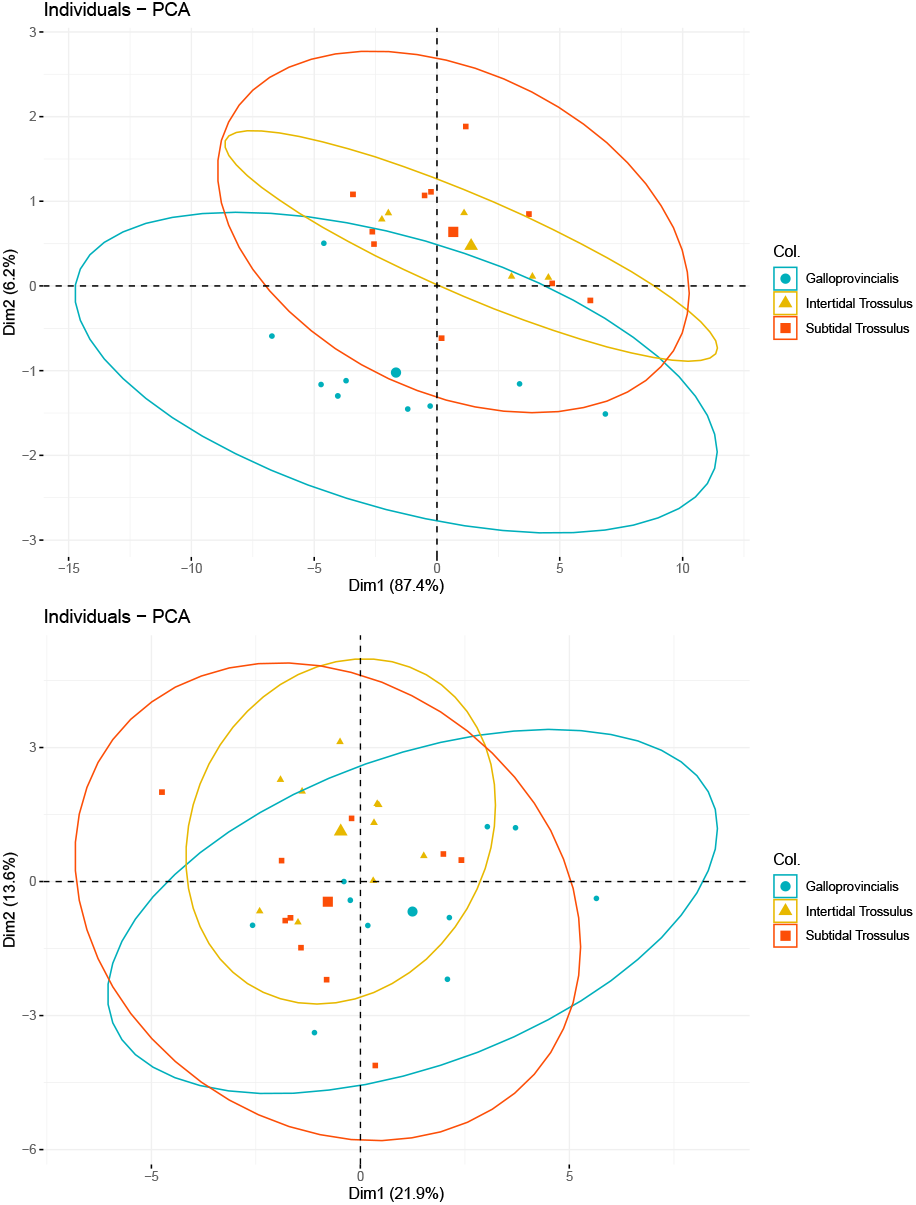
PCA plot based on (1) ^1^H NMR identified metabolites and (2) standard amino acids in intertidal *Mytilus trossulus*, subtidal *M. trossulus* and subtidal *M. galloprovincialis* after subjection to control (pH = 7.9) and low (pH = 7.5) pH treatment (n = 5).

**Figure S2:**
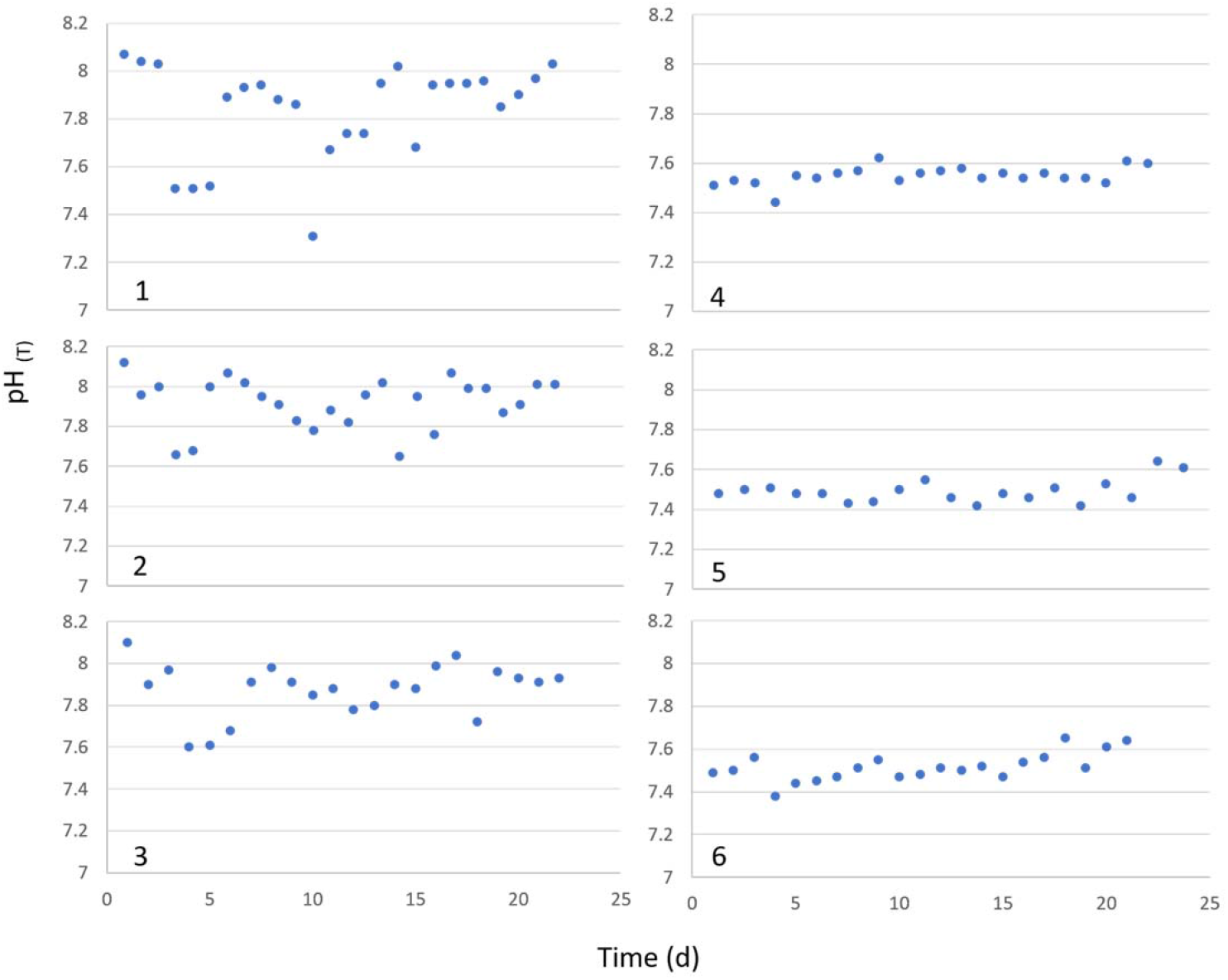
pH measurements from each incubator during the observation period. Incubators 1-3 were set to control conditions (pH = 7.9), while incubators 4-6 were set to acidified conditions (pH = 7.5).

**Table S1:** Carbonate measurement data from the burke-o-lator system at Hakai Institute. Bold numbers indicate anomalous values.

| Treatment | Incubator | DIC<br>($\mu\text{mol/kg}$) | TA<br>($\mu\text{mol/kg}$) | pCO<sub>2</sub><br>($\mu\text{atm}$) | pH<br>(total) | Aragonite<br>($\Omega$) | Calcite<br>($\Omega$) |
| --- | --- | --- | --- | --- | --- | --- | --- |
| Control<br>(start) | 3 | 1230.64 | 1335.34 | 176.67 | 8.19 | 1.07 | 1.76 |
| Control<br>(start) | 1 | 1260.26 | 1362.46 | 188.24 | 8.17 | 1.06 | 1.74 |
| Control<br>(start) | 2 | 1272.46 | 1316.33 | 361.12 | 7.90 | 0.58 | 0.96 |
| **Control**<br>**(end)** | 3 | **916.80** | **986.24** | **170.48** | 8.08 | 0.63 | 1.04 |
| **Control**<br>**(end)** | 2 | **705.92** | **776.96** | **115.85** | 8.13 | 0.55 | 0.90 |
| **Control**<br>**(end)** | 1 | **798.98** | **883.86** | **118.12** | 8.17 | 0.69 | 1.13 |
| Acidified<br>(start) | 5 | 1421.88 | 1433.95 | 621.42 | 7.71 | 0.42 | 0.70 |
| Acidified<br>(start) | 6 | 1516.80 | 1455.56 | 1548.91 | 7.34 | 0.19 | 0.31 |
| Acidified<br>(start) | 4 | 1599.63 | 1607.00 | 735.82 | 7.69 | 0.45 | 0.75 |
| Acidified<br>(end) | 5 | 1620.13 | 1637.31 | 678.28 | 7.73 | 0.52 | 0.85 |
| Acidified<br>(end) | 4 | 1712.32 | 1653.04 | 1600.13 | 7.37 | 0.24 | 0.39 |
| Acidified<br>(end) | 6 | 1380.71 | 1357.44 | 985.94 | 7.49 | 0.26 | 0.42 |

**Table S2:** Mean percentage (± standard deviation) of all fatty acids after subjection to control (pH = 7.9) and acidified (pH = 7.5) conditions (n = 5). The summarized degree of SFA, MUFA and PUFA is presented.

| Fatty acid | *M. trossulus* - intertidal |  | *M. trossulus* - subtidal |  | *M. galloprovincialis* - subtidal |  |
| --- | --- | --- | --- | --- | --- | --- |
|  | Acidified | Control | Acidified | Control | Acidified | Control |
| 14:0 | 0.40±0.07 | 0.4±0.09 | 0.54±0.24 | 0.3±0.05 | 0.62±0.14 | 0.37±0.07 |
| 15:0 | 0.52±0.04 | 0.51±0.05 | 0.69±0.21 | 0.41±0.05 | 0.63±0.26 | 0.44±0.09 |
| 15:0iso | 0.07±0.06 | 0.09±0.1 | 0.2±0.19 | 0.16±0.06 | 0.00 | 0.08±0.05 |
| 16:0 | 9.53±0.89 | 8.72±1.21 | 16.56±6.62 | 10.21±1.21 | 12.95±5.17 | 8.83 ±1.44 |
| 17:0 | 0.45±0.05 | 0.43±0.04 | 0.88±0.58 | 0.38±0.05 | 0.61±0.28 | 0.41±0.09 |
| 17:0iso | 0.3±0.07 | 0.29±0.05 | 0.29±0.13 | 0.12±0.02 | 0.24±0.07 | 0.19±0.02 |
| 17:0ante | 0.17±0.03 | 0.14±0.02 | 0.19±0.09 | 0.08±0.01 | 0.17±0.06 | 0.12±0.03 |
| 18:0 | 2.57±0.34 | 2.53±0.17 | 4.30±1.98 | 2.28±0.38 | 3.38±1.26 | 2.53±0.67 |
| 18:0iso | 0.11±0.01 | 0.1±0.02 | 0.21±0.11 | 0.08±0.01 | 0.21±0.07 | 0.17±0.03 |
| **ΣSFA** | **14.12±1.31** | **13.2±1.27** | **23.86±9.57** | **14.03±1.53** | **18.81±7.26** | **13.12±2.3** |
| 16:1n7 | 1.01±0.1 | 1.2±0.22 | 1.58±0.33 | 0.97±0.19 | 1.71±0.33 | 1.07±0.27 |
| 17:1n7 | 1.71±0.63 | 4.07±4.16 | 1.94±1.14 | 1.43±1.1 | 2.26±1.63 | 2.04±2.15 |
| 18:1n7 | 1.48±0.12 | 1.37±0.37 | 1.91±0.52 | 1.21±0.2 | 2.08±0.58 | 1.5±0.34 |
| 18:1n9 | 1.04±0.16 | 1.01±0.24 | 1.23±0.32 | 1.07±0.29 | 1.2±0.33 | 0.97±0.26 |
| 20:1n7 | 0.65±0.1 | 0.60±0.14 | 0.75±0.2 | 0.34±0.06 | 0.97±0.27 | 0.61±0.12 |
| 20:1n9 | 4.31±0.3 | 4.12±0.46 | 6.21±1.13 | 4.86±0.61 | 5.43±1.32 | 4.37±0.48 |
| 20:1n11 | 0.67±0.14 | 0.74±0.19 | 0.74±0.15 | 0.53±0.08 | 1.14±0.19 | 0.82±0.15 |
| **ΣMUFA** | **10.87±1.1** | **13.13±3.68** | **14.36±3.33** | **10.41±1.02** | **14.78±1.65** | **11.39±1.63** |
| 18:2n6 | 0.91±0.1 | 0.94±0.35 | 1.46±0.67 | 1.31±0.25 | 0.79±0.36 | 0.64±0.14 |
| 18:3n3 | 1.12±0.07 | 1.21±0.5 | 1.04±0.57 | 1.52±0.19 | 0.31±0.1 | 0.36±0.09 |
| 18:4n3 | 0.76±0.19 | 1.07±0.41 | 0.67±0.25 | 1.09±0.28 | 0.39±0.13 | 0.33±0.11 |
| 20:2n6 | 0.44±0.09 | 0.46±0.18 | 0.51±0.22 | 0.67±0.08 | 0.32±0.09 | 0.3±0.04 |
| 20:2n | 9.41±0.76 | 8.82±1.05 | 11.96±1.21 | 9.87±0.58 | 9.78±1.79 | 8.67±1.41 |
| 20:3n6 | 0.11±0.02 | 0.12±0.04 | 0.00 | 0.09±0.02 | 0.16±0.03 | 0.12±0.02 |
| 20:4n6 | 9.7±1.1 | 9.81±0.9 | 3.76±2.61 | 6.07±0.76 | 6.95±2.43 | 9.7±1.37 |
| 20:5n3 | 9.78±1.15 | 10.74±1.97 | 4.87±3.59 | 12.19±1.05 | 7.0±2.98 | 10.21±1.68 |
| 22:2NMI | 11.54±0.84 | 10.86±1.5 | 12.72±0.95 | 9.55±0.66 | 13.53±1.5 | 12.19±1.14 |
| 22:6n3 | 14.13±0.99 | 13.73±0.69 | 6.36±4.86 | 16.22±0.48 | 9.7±4.61 | 15.88±2.06 |
| TMTD | 0.14±0.19 | 0.1±0.1 | 0.51±0.52 | 0.09±0.07 | 0.17±0.12 | 0.12±0.11 |
| PUFA0 | 1.19±0.23 | 0.98±0.1 | 1.51±0.7 | 2.36±0.42 | 0.72±0.07 | 0.81±0.17 |
| PUFA1 | 2.62±0.52 | 2.47±0.25 | 2.84±1.49 | 4.06±0.34 | 1.37±0.42 | 1.72±0.42 |
| PUFA2 | 0.93±0.26 | 0.86±0.23 | 0.4±0.11 | 0.38±0.08 | 0.91±0.33 | 1.09±0.12 |
| PUFA3 | 0.68±0.11 | 0.64±0.07 | 0.46±0.1 | 0.52±0.05 | 0.77±0.18 | 1.14±0.22 |
| PUFA4 | 11.54±0.8 | 10.86±1.5 | 12.72±0.95 | 9.55±0.66 | 13.52±1.5 | 12.19±1.14 |
| **ΣPUFA** | **75.0±2.05** | **73.67±2.93** | **61.78±12.27** | **75.56±1.65** | **66.4±8.57** | **75.49±2.1** |

**Table S3:** Estimated regression parameters, standard errors, z-values and p-values for the binomial generalized linear models (GLM).

| GLM Model<br>(Binomial distribution) | Estimate | s.e | z-value | p-value |
| --- | --- | --- | --- | --- |
| Subtidal *Mytilus galloprovincialis* (Explained deviance = 95.37%) |  |  |  |  |
| Intercept | 11.96 | 1.81 | 6.59 | < 0.0001 |
| Air temperature | 1.58 | 0.24 | 6.63 | < 0.0001 |
| Low pH | -1.42 | 0.50 | -2.82 | < 0.0001 |
| Subtidal *Mytilus trossulus* (Explained deviance = 91.49%) |  |  |  |  |
| Intercept | 8.11 | 1.13 | 7.20 | < 0.0001 |
| Air temperature | 0.88 | 0.13 | 6.84 | < 0.0001 |
| Low pH | -1.31 | 0.43 | -3.05 | < 0.0001 |
| Intertidal *Mytilus trossulus* (Explained deviance = 93.94%) |  |  |  |  |
| Intercept | 12.71 | 1.83 | 6.93 | < 0.0001 |
| Air temperature | 1.21 | 0.17 | 6.79 | < 0.0001 |
| Low pH | -3.29 | 0.70 | -4.73 | < 0.0001 |
| Survival among mussel categories (Explained deviance = 90.7%) |  |  |  |  |
| Intercept | 7.08 | 0.79 | 11.15 | < 0.0001 |
| Air temperature | 0.99 | 0.10 | 11.32 | < 0.0001 |
| Intertidal *Mytilus trossulus* | 1.80 | 0.34 | 5.97 | < 0.0001 |
| Subtidal *Mytilus trossulus* | 1.25 | 0.32 | 4.45 | < 0.0001 |

**Table S4:**
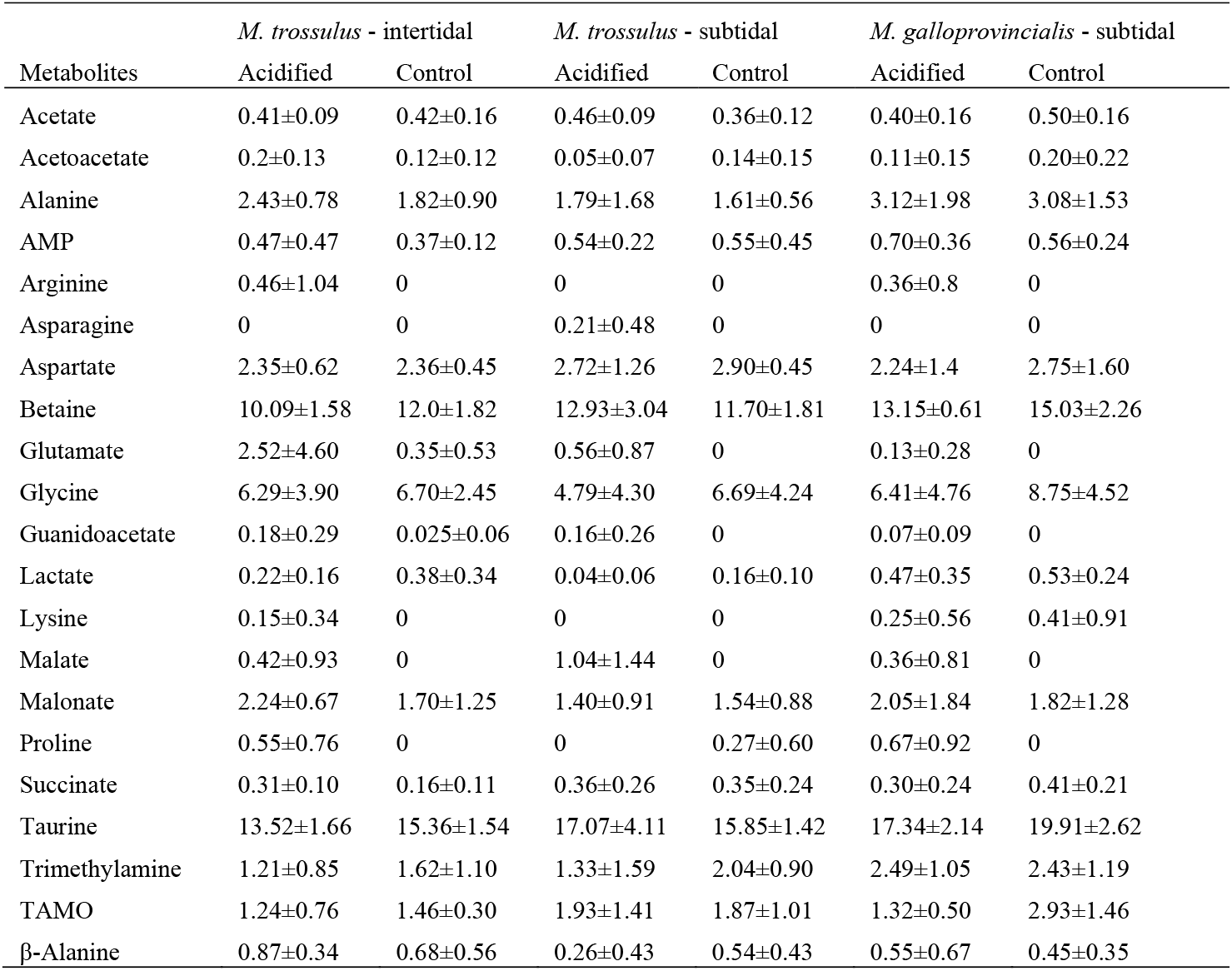
Mean (± standard deviation) metabolite concentration (nmol / 100 g ww gill tissue) detected from the H^1^NMR analysis after subjection to control (pH = 7.9) and acidified (pH = 7.5) conditions (n = 5).

| Metabolites | *M. trossulus* - intertidal |  | *M. trossulus* - subtidal |  | *M. galloprovincialis* - subtidal |  |
| --- | --- | --- | --- | --- | --- | --- |
|  | Acidified | Control | Acidified | Control | Acidified | Control |
| Acetate | 0.41±0.09 | 0.42±0.16 | 0.46±0.09 | 0.36±0.12 | 0.40±0.16 | 0.50±0.16 |
| Acetoacetate | 0.2±0.13 | 0.12±0.12 | 0.05±0.07 | 0.14±0.15 | 0.11±0.15 | 0.20±0.22 |
| Alanine | 2.43±0.78 | 1.82±0.90 | 1.79±1.68 | 1.61±0.56 | 3.12±1.98 | 3.08±1.53 |
| AMP | 0.47±0.47 | 0.37±0.12 | 0.54±0.22 | 0.55±0.45 | 0.70±0.36 | 0.56±0.24 |
| Arginine | 0.46±1.04 | 0 | 0 | 0 | 0.36±0.8 | 0 |
| Asparagine | 0 | 0 | 0.21±0.48 | 0 | 0 | 0 |
| Aspartate | 2.35±0.62 | 2.36±0.45 | 2.72±1.26 | 2.90±0.45 | 2.24±1.4 | 2.75±1.60 |
| Betaine | 10.09±1.58 | 12.0±1.82 | 12.93±3.04 | 11.70±1.81 | 13.15±0.61 | 15.03±2.26 |
| Glutamate | 2.52±4.60 | 0.35±0.53 | 0.56±0.87 | 0 | 0.13±0.28 | 0 |
| Glycine | 6.29±3.90 | 6.70±2.45 | 4.79±4.30 | 6.69±4.24 | 6.41±4.76 | 8.75±4.52 |
| Guanidoacetate | 0.18±0.29 | 0.025±0.06 | 0.16±0.26 | 0 | 0.07±0.09 | 0 |
| Lactate | 0.22±0.16 | 0.38±0.34 | 0.04±0.06 | 0.16±0.10 | 0.47±0.35 | 0.53±0.24 |
| Lysine | 0.15±0.34 | 0 | 0 | 0 | 0.25±0.56 | 0.41±0.91 |
| Malate | 0.42±0.93 | 0 | 1.04±1.44 | 0 | 0.36±0.81 | 0 |
| Malonate | 2.24±0.67 | 1.70±1.25 | 1.40±0.91 | 1.54±0.88 | 2.05±1.84 | 1.82±1.28 |
| Proline | 0.55±0.76 | 0 | 0 | 0.27±0.60 | 0.67±0.92 | 0 |
| Succinate | 0.31±0.10 | 0.16±0.11 | 0.36±0.26 | 0.35±0.24 | 0.30±0.24 | 0.41±0.21 |
| Taurine | 13.52±1.66 | 15.36±1.54 | 17.07±4.11 | 15.85±1.42 | 17.34±2.14 | 19.91±2.62 |
| Trimethylamine | 1.21±0.85 | 1.62±1.10 | 1.33±1.59 | 2.04±0.90 | 2.49±1.05 | 2.43±1.19 |
| TAMO | 1.24±0.76 | 1.46±0.30 | 1.93±1.41 | 1.87±1.01 | 1.32±0.50 | 2.93±1.46 |
| β-Alanine | 0.87±0.34 | 0.68±0.56 | 0.26±0.43 | 0.54±0.43 | 0.55±0.67 | 0.45±0.35 |

**Table S5:**
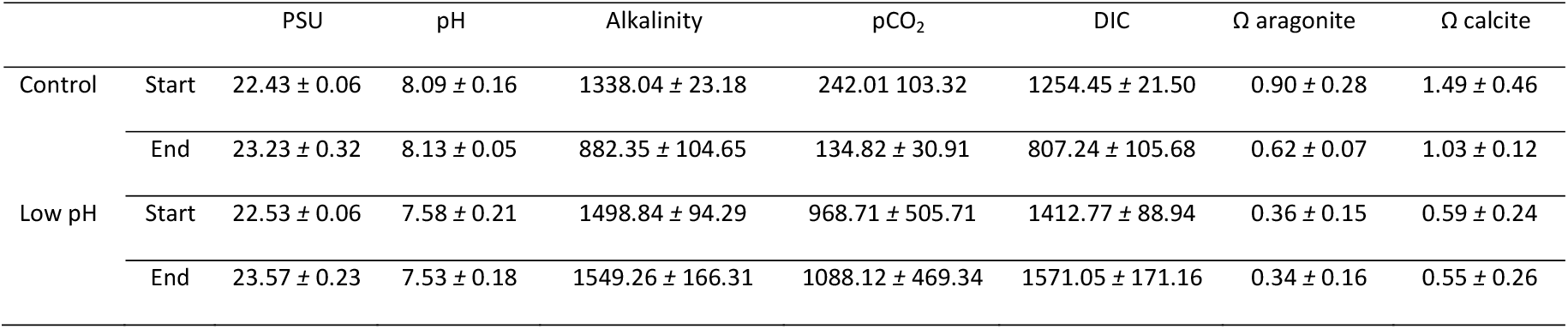
Mean (± standard deviation) carbonate measurements taken from discrete water samples at the start and end of the exposure period.

